# Role of microRNA-21 in hypertrophic cardiac remodeling

**DOI:** 10.1101/850156

**Authors:** Ken Watanabe, Taro Narumi, Tetsu Watanabe, Yoichiro Otaki, Tetsuya Takahashi, Tomonori Aono, Jun Goto, Taku Toshima, Takayuki Sugai, Masahiro Wanezaki, Daisuke Kutsuzawa, Shigehiko Kato, Harutoshi Tamura, Satoshi Nishiyama, Hiroki Takahashi, Takanori Arimoto, Tetsuro Shishido, Masafumi Watanabe

## Abstract

Hypertension is a major public health problem among with aging population worldwide. It causes cardiac remodeling, including hypertrophy and interstitial fibrosis, which leads to development of hypertensive heart disease (HHD). Although microRNA-21 (miR-21) is associated with fibrogenesis in multiple organs, its impact on hypertrophic cardiac remodeling in hypertension is not known. Circulating miR-21 level was higher in patients with HHD than that in the control subjects. It also positively correlated with serum myocardial fibrotic markers. MiR-21 expression levels were significantly upregulated in the mice hearts after angiotensin II (Ang II) infusion or transverse aortic constriction (TAC) compared with control mice. Expression level of programmed cell death 4 (PDCD4), a main target of miR-21, was significantly decreased in Ang II infused mice and TAC mice compared with control mice. Expression levels of transcriptional activator protein 1 (AP-1) and transforming growth factor-β1 (TGF-β1), which were downstream targets of PDCD4, were increased in Ang II infused mice and TAC mice compared with control mice. *In vitro*, mirVana-miR-21-specific inhibitor attenuated Ang II-induced PDCD4 downregulation and contributed to subsequent deactivation of AP-1/TGF-β1 signaling pathway in neonatal rat cardiomyocytes. Thus, suppression of miR-21 prevents hypertrophic cardiac remodeling by regulating PDCD4, AP-1, and TGF-β1 signaling pathway.

## Introduction

Hypertension is a major public health concern among the elderly population worldwide. It is associated with increased risk of adverse cardiovascular events [1]. An epidemiological report indicated that 31.1 % of adults in the world (1.39 billion people) had hypertension in 2010 [2]. Hypertension increases the risk of developing hypertension-induced organ damages, such as hypertensive heart disease (HHD), hypertensive encephalopathy, and nephrosclerosis [3]. HHD is one of the most important hypertension-induced organ damages [4]. According to the Framingham Heart Study, 20 mmHg increase in systolic blood pressure contributes to 56% increased risk for heart failure [5]. Furthermore, it was reported that HHD is a common pathophysiology of heart failure with preserved ejection fraction [6, 7]. Hypertension causes cardiac remodeling characterized by cardiac fibrosis, which contributes to progression of heart failure [3, 4].

MicroRNAs (miRs) are small non-coding RNAs that regulate post-transcriptional gene expressions. They have been shown to play an important role in fibrogenic process in multiple organs [8]. In the present study, we focused on the fibrogenic function of miR-21, which is a ubiquitously expressed miR that is reported to have a pivotal role in development of tissue fibrosis [9]. Transforming growth factor-β1 (TGF-β1), a pleiotropic and multifactorial cytokine involved in many biological processes, plays a crucial role in the pathogenesis of cardiac remodeling in hypertension [10]. It has been demonstrated that miR-21 can promote TGF-β1 signaling [11, 12]. On the other hand, miR-21 has been found to be upregulated by TGF-β1 [13]. This interrelationship forms a positive feedback loop, which may exacerbate the fibrogenic process. Previous studies have also demonstrated the contribution of miR-21 in patients with aortic stenosis, hypertrophic cardiomyopathy, and dilated cardiomyopathy [14–16]. However, the impact of miR-21 on the pathogenesis of hypertrophic cardiac remodeling in hypertension is still not clear.

We hypothesized that miR-21 deteriorates hypertrophic cardiac remodeling by enhancing TGF-β1 signaling pathway through suppressing its target gene expression. In the present study, we investigated the following: (1) miR-21 expression levels in patients with HHD; (2) miR-21 expression levels and its downstream signaling in animal model of hypertrophic cardiac remodeling by transverse aortic constriction (TAC) or angiotensin II (Ang II) infusion; (3) the function of miR-21 in cardiac remodeling process in response to Ang II stimulation *in vitro*; (4) the therapeutic potential of miR-21 inhibitor in hypertrophic cardiac remodeling *in vitro*.

## Materials and Methods

### Human studies

The present study included 10 patients with HHD and 10 control patients who were assessed to rule out cardiomyopathy and heart failure, and had normal cardiac function (Table 1). Endomyocardial biopsies (EMBs) were collected from the patients who had left ventricular hypertrophy and suspected some types of cardiomyopathy. EMBs were taken from left ventricle with a total of 4 to 6 samples through the femoral arteries. EMBs were analyzed in 3 HHD patients who were excluded other cardiomyopathy based on EMBs and other clinical data, and 3 control patients who had transient left ventricular dysfunction and suspected myocarditis but were eventually ruled out cardiomyopathy. The final diagnosis of HHD were made by two expert cardiologists based on angiography, echocardiographic data, clinical background, and medical history. Written informed consent was obtained from all patients before the study. The protocol was performed in accordance to the Helsinki Declaration and was approved by the human investigations committee of Yamagata University School of Medicine.

**Table 1.** Clinical characteristics of 10 control subjects and 10 HHD patients.

| Variables | Control patients<br>*n* = 10 | HHD patients<br>*n* = 10 | *P* value |
| --- | --- | --- | --- |
| Age (years old) | 61 ± 8 | 58 ± 12 | ns |
| Male / Female | 6 / 4 | 7 / 3 | ns |
| BMI (kg/m<sup>2</sup>) | 23.7 ± 1.9 | 26.1 ± 5.9 | ns |
| Hypertension, n (%) | 3 (30) | 10 (100) | <0.05 |
| Diabetes mellitus, n (%) | 1 (10) | 6 (60) | <0.05 |
| Dyslipidemia, n (%) | 3 (30) | 4 (40) | ns |
| NYHA functional class III-IV, n (%) | 0 (0) | 4 (40) | <0.05 |
| Echocardiographic data |  |  |  |
| LVEDD (mm) | 47 ± 5 | 55 ± 9 | <0.05 |
| LVEF (%) | 68 ± 6 | 52 ± 14 | <0.05 |
| IVSD (mm) | 9 ± 2 | 14 ± 2 | <0.05 |
| LVPWD (mm) | 10 ± 1 | 13 ± 2 | <0.05 |
| Blood examination |  |  |  |
| eGFR (mL/min/1.73 m<sup>2</sup>) | 92.0 ± 18.9 | 56.3 ± 25.5 | <0.05 |
| BNP (pg/mL) | 36 (17 - 70) | 463 (143 - 713) | <0.05 |
| Medications |  |  |  |
| ACEIs and/or ARBs, n (%) | 3 (30) | 10 (100) | <0.05 |
| CCBs, n (%) | 1 (10) | 7 (70) | <0.05 |
| Diuretics, n (%) | 0 (0) | 6 (60) | <0.05 |
| Statins, n (%) | 3 (30) | 5 (50) | ns |

Data are expressed as mean ± SD, number (percentage), or median (interquartile range). ACEIs, angiotensin-converting enzyme inhibitors; ARBs, angiotensin II receptor blockers; BMI, body mass index; BNP, B-type natriuretic peptide; CCBs, calcium- channel blockers; eGFR, estimated glomerular filtration rate; HHD, hypertensive heart disease; IVSD, interventricular septum diameter; LVEDD, left ventricular end-diastolic diameter; LVEF, left ventricular ejection fraction; LVPWD, left ventricular posterior wall diameter; NYHA, New York Heart Association.

### Measurement of circulating miR-21 levels and biochemical assays

Blood samples were collected in the early morning within 24 hours after admission, centrifuged at 3000g for 15 min at 4 °C, and the obtained serum was stored at −80 °C. Circulating miRs were isolated from 300 µL serum by using a NucleoSpin microRNA isolation kit (TaKaRa, Otsu, Japan).

Serum carboxy-terminal telopeptide of type I collagen (I-CTP) concentrations were determined by radioimmunoassay (Orion Diagnostica, Finland) [17]. Serum procollagen type III N-terminal propeptide (P3NP) levels were measured with enzyme-linked immunosorbent assay (ELISA) kit (MyBioSource, San Diego, CA, USA).

### Animal treatment regimens

Hypertension-induced cardiac remodeling models were established by Ang II infusions or TAC surgery [18, 19]. Briefly, Ang II was infused with ALZET osmotic pumps (1.5 mg/kg/day) as we previously described [18]. Cardiac function, dimension, and blood pressure were assessed after 2 weeks from Ang II infusion. The mice were sacrificed by intraperitoneal injection of a combination of ketamine (1g/kg) and xylazine (100 mg/kg), and the heart samples were obtained for the biochemical and histopathological study. TAC surgery was performed to induce chronic pressure overload as we previously described [20]. Briefly, 8- to 10-week-old mice were anesthetized by intraperitoneal injection with a mixture of ketamine (80 mg/kg) and xylazine (8 mg/kg). Animals were intubated and ventilated with a rodent ventilator (Harvard Apparatus, Holliston, MA, USA). The transverse aortic arch was ligated (7-0 prolene) between the right innominate and left common carotid arteries with a 27-gauge needle, and then the needle was promptly removed leaving a discrete region of stenosis. Cardiac remodeling was assessed after 4 weeks from surgery. All experimental procedures were performed according to the animal welfare regulations of Yamagata University School of Medicine, and the study protocol was approved by the Animal Subjects Committee of Yamagata University School of Medicine. The investigation conformed to the Guide for the Care and Use of Laboratory Animals published by the US National Institutes of Health (NIH Publication, 8th Edition, 2011).

### Neonatal rat cardiomyocyte isolation, cell culture, and treatment

Isolation and culture of neonatal rat cardiomyocytes (NRCMs) were performed as we previously described [21]. Briefly, ventricles were obtained from 1- to 2-day-old Sprague-Dawley rat pups after euthanasia by decapitation, and cardiomyocytes were isolated by digestion with collagenase. Cardiomyocytes were kept in serum-supplemented (10% fatal bovine serum, FBS) Dulbecco’s Modified Eagle Medium (DMEM, Thermo Fisher Scientific, MA, USA). Primary culture of cardiofibroblasts were obtained as previously described [22]. Briefly, ventricles of Sprague-Dawley rat pups were digested with collagenase, and resuspended in DMEM with 10% FBS. Cells were then seeded into 10-cm culture dishes and cultured at 37 °C for 2h. Unattached cells were discarded, and attached cells were cultured in DMEM with 10% FBS. NRCMs were transfected with small interfering RNA (siRNA) specific for programmed cell death 4 (PDCD4) (Thermo Fisher Scientific), 10-nM mirVana hsa-miR-21 specific inhibitor (Thermo Fisher Scientific), or mirVana miRNA inhibitor Negative Control (Thermo Fisher Scientific) using Lipofectamine 3000 Reagent (Thermo Fisher Scientific) according to the manufacturer’s instructions. The medium was replaced with DMEM with 10% FBS after transfection for 4h. NRCMs were stimulated with 1 µM Ang II for 24 hours of serum starvation.

### Western blotting

The total protein extracts were prepared with radio-immunoprecipitation assay (RIPA) buffer as we previously reported [21]. Equal amounts of protein were subjected to 10% SDS-PAGE and transferred to polyvinylidene difluoride membranes. The membranes were probed overnight at 4 °C with the following primary antibodies: PDCD4 (Santa Cruz, Dallas, TX, USA, sc-376430), c-Jun (Cell Signaling Technology, Danvers, MA, USA, #9165), TGF-β1 (Cell Signaling Technology, #3711), phospho-transforming growth factor-β-activated kinase 1 (p-TAK1) (Cell Signaling Technology, #4537), TAK1 (Cell Signaling Technology, #4505), β-tubulin (Cell Signaling Technology, #2146). Protein expression levels were normalized to that of β-tubulin.

### RNA extraction and quantitative reverse transcription polymerase chain reaction (qRT-PCR)

Total RNA was isolated from human endomyocardial biopsy specimens, mouse whole heart, and NRCMs using TRIzol reagent (Thermo Fisher Scientific) as we previously described [23]. For miRs screening assay, first strand cDNA of miRs was synthesized, and PCR reaction was performed using a miR-X miRNA qRT-PCR SYBR Kit (TaKaRa) according to the manufacturer’s instructions. For other studies, first strand cDNA was synthesized using a Superscript IV First-strand cDNA synthesis kit (Thermo Fisher Scientific) and quantitative RT-PCR (qRT-PCR) was performed with SYBR Green Real-Time PCR Master Mixes (Thermo Fisher Scientific) according to the manufacturer’s instructions. Gene expressions was normalized to U6 for miR assay and β-actin for other assays.

### Histopathological examinations

Biopsy samples of human cardiomyocyte and mice heart samples were fixed with 4% formalin and embedded in paraffin. Sections of 3–5 µm thickness were stained with hematoxylin-eosin (HE) or Massons’s trichrome stain for histopathological analysis as we previously described [20]. The extent of myocardial interstitial fibrosis was evaluated using a microscope and attached software (BZ-X710; Keyence, Osaka, Japan).

### Statistical analysis

All values are expressed as mean ± standard error of mean (SEM). Statistical differences among groups were evaluated with one-way analysis of variance (ANOVA) followed by Tukey-Kramer post hoc tests. Correlations between the circulating miRs levels and biomarkers of cardiac fibrosis were analyzed by using Pearson’s correlation coefficient. A value of *P* < 0.05 was considered statistically significant. All statistical analyses were performed with a standard software package (JMP version 12; SAS institute, Cary, NC, USA).

## Results

### MiRs expression levels in patients with hypertensive heart disease

To investigate the expression levels of fibrosis-associated miRs according to previous report [24], we first measured the levels of circulating miR-21, miR-29, miR-30, and miR-133 in patients with HHD. Circulating miR-21 levels were significantly increased in patients with HHD compared with those of control subjects. On the other hand, circulating miR-29, miR-30, and miR-133 levels were significantly decreased in patients with HHD (Fig. 1A). HE and Masson’s trichrome staining revealed that significant cardiac hypertrophy and fibrosis was observed in the heart section from patients with HHD (Fig. 1B). MiR-21 levels were significantly increased in the heart samples of patients with HHD compared with those of the normal subjects. In contrast, miR-29, miR-30, and miR-133 levels tended to be decreased in patients with HHD, but the differences were not statistically significant (Fig. 1C). We measured serum I-CTP and P3NP levels as markers of myocardial fibrosis [17, 25]. Serum I-CTP and P3NP levels were significantly higher in patients with HHD compared with those of control subjects (Fig. 1D). As shown in Fig. 1E, there were significant positive correlations between circulating miR-21 levels and serum I-CTP (R = 0.560) and P3NP (R = 0.477). However, there were no significant correlations between other miRs and I-CTP (miR-29: R = −0.215; miR-30: R = −0.068; miR-133: R = 0.268) and P3NP (miR-29: R = −0.302; miR-30: R = −0.263; miR-133: R = −0.138) levels.

**Fig 1.**
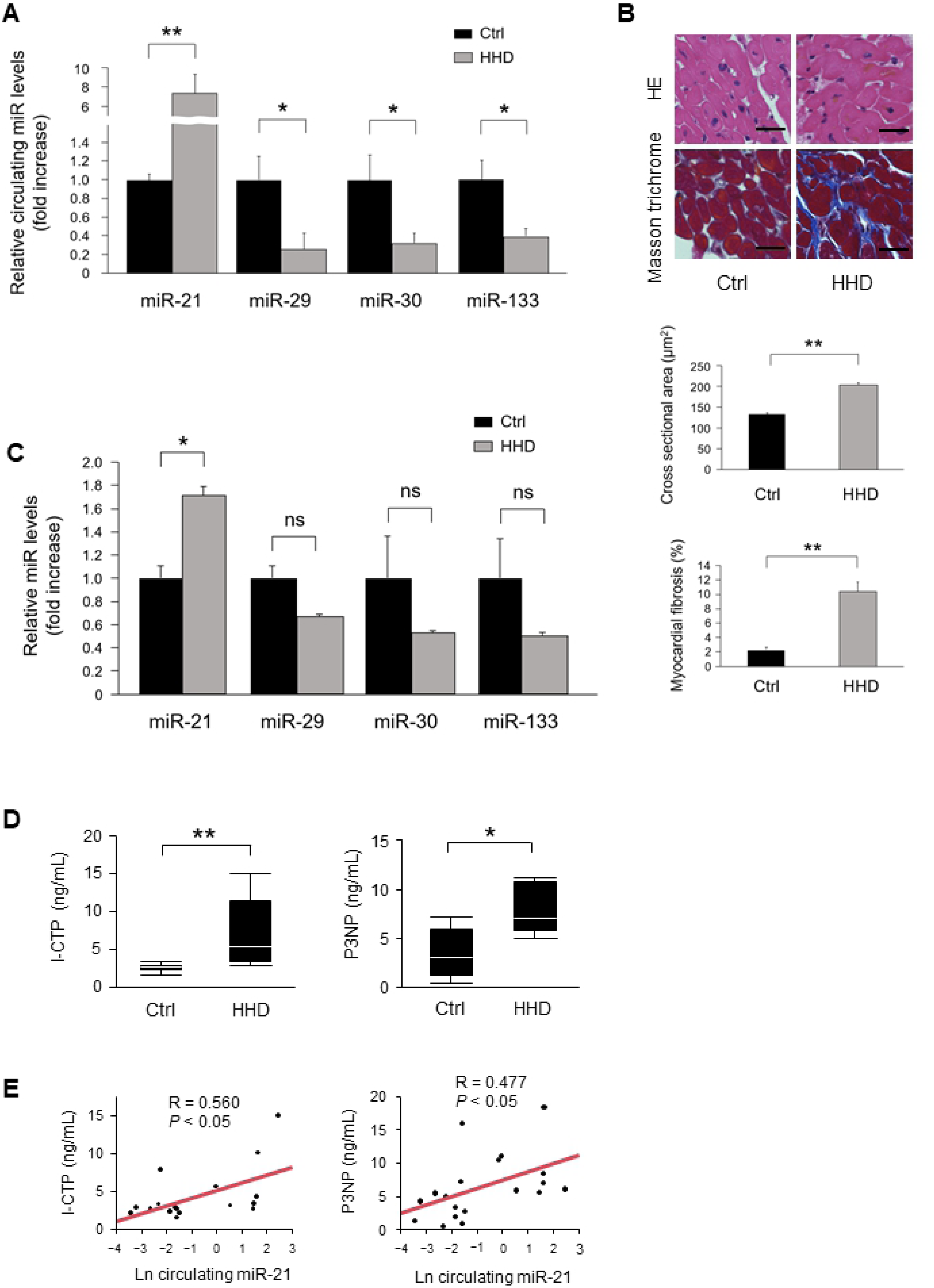
Association between miRs expressions and cardiac remodeling in patients with HHD. **(A)** Circulating miRs expressions in patients with HHD (n = 10 per group). **(B)** Representative images and analysis of cardiac remodeling by HE and Masson’s trichrome staining in heart samples from HHD patients and normal subjects (n = 3 per group). Scale bars = 20 µm. **(C)** Expression of miRs levels in heart samples from patients with HHD (n = 3 per group). **(D)** Serum I-CTP and P3NP levels in patients with HHD. **(E)** Circulating miR-21 levels were positively correlated with serum I-CTP and P3NP levels in patients with HHD. Data are expressed as mean ± SEM. \**P* < 0.05, \*\**P* < 0.01.

### MiR-21 expression levels in Ang II infused mice and TAC mice

MiR-21 expression levels were significantly increased in Ang II infused mice hearts compared with those of sham mice (Fig. 2A). Cardiac remodeling was detected by HE and Masson’s trichrome staining in Ang II infused mice hearts but not in sham mice (Fig. 2B). Alpha smooth muscle actin (α-SMA) mRNA expression was significantly upregulated in the Ang II infused mice hearts compared with those of the sham mice (Fig. 2C). Similarly, miR-21 expression levels were significantly increased in the heart of TAC mice compared with those of sham-operated mice (Fig. 2D). Cardiac remodeling was also detected by HE and Masson’s trichrome staining in TAC mice but not in the sham-operated mice (Fig. 2E). α-SMA mRNA expression was significantly upregulated in the TAC mice hearts compared with those of the sham-operated mice (Fig. 2F). Echocardiographic and hemodynamic data of Ang II infused mice and TAC operated mice are shown in Table 2.

**Fig 2.**
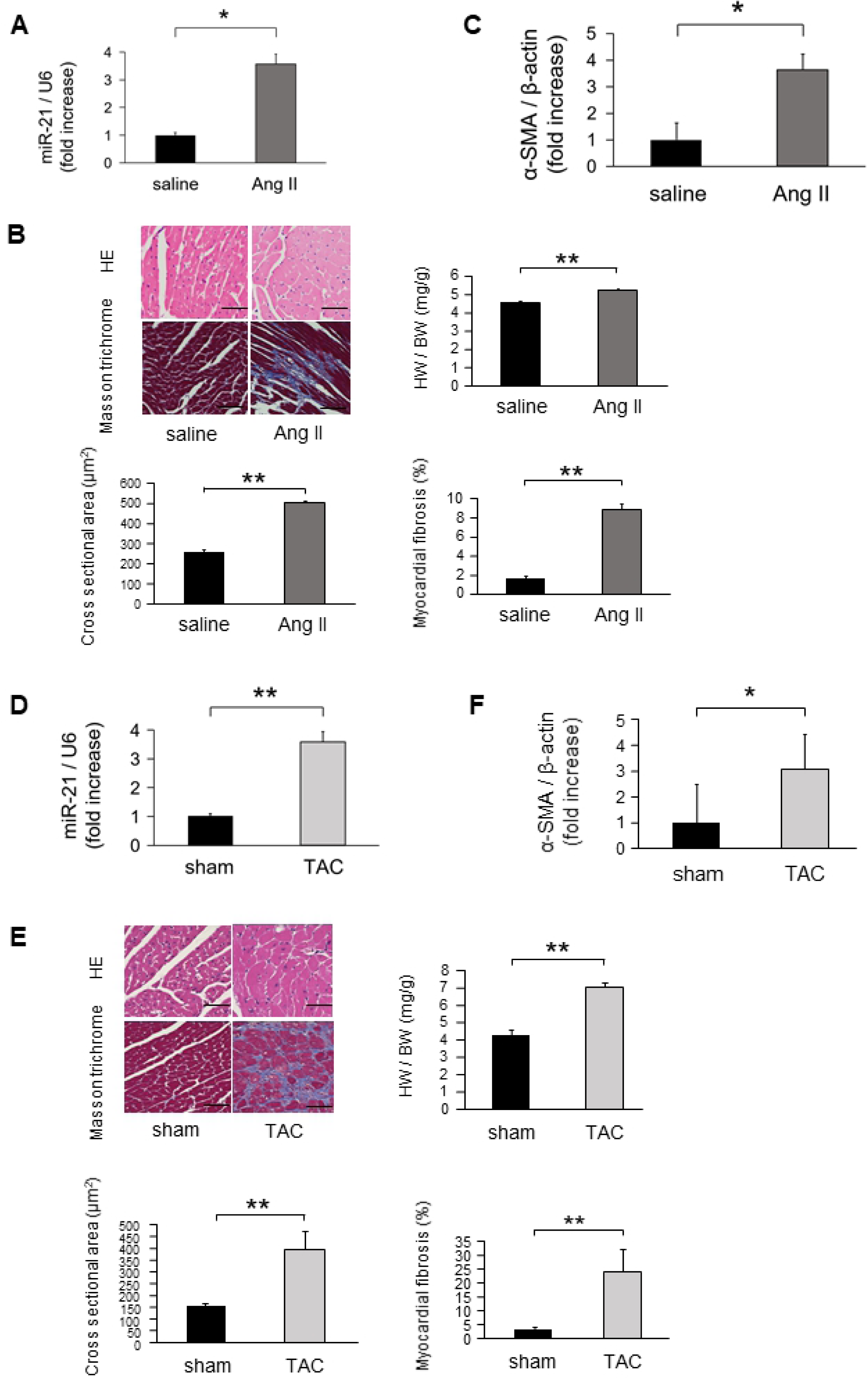
The crucial role of miR-21 in cardiac remodeling in Ang II infused and TAC mice models. **(A)** MiR-21 expression levels in Ang II infused mice (n = 6 per group). **(B)** Representative images and analysis of cardiac remodeling by HE and Masson’s trichrome staining in left ventricular sections in Ang II infused mice hearts (n = 6 per group). Scale bars = 50 µm. **(C)** α-SMA expression in Ang II infused mice (n = 6 per group). **(D)** MiR-21 expression levels in the heart samples of TAC-operated mice (n = 6 per group). **(E)** Representative images and analysis of cardiac remodeling by HE and Masson’s trichrome staining in left ventricular sections of the TAC-operated mice hearts (n = 6 per group). Scale bars = 50 µm. **(F)** α-SMA expression in TAC-operated mice (n = 6 per group). Data are expressed as mean ± SEM. \**P* < 0.05, \*\**P* < 0.01.

**Table 2.**
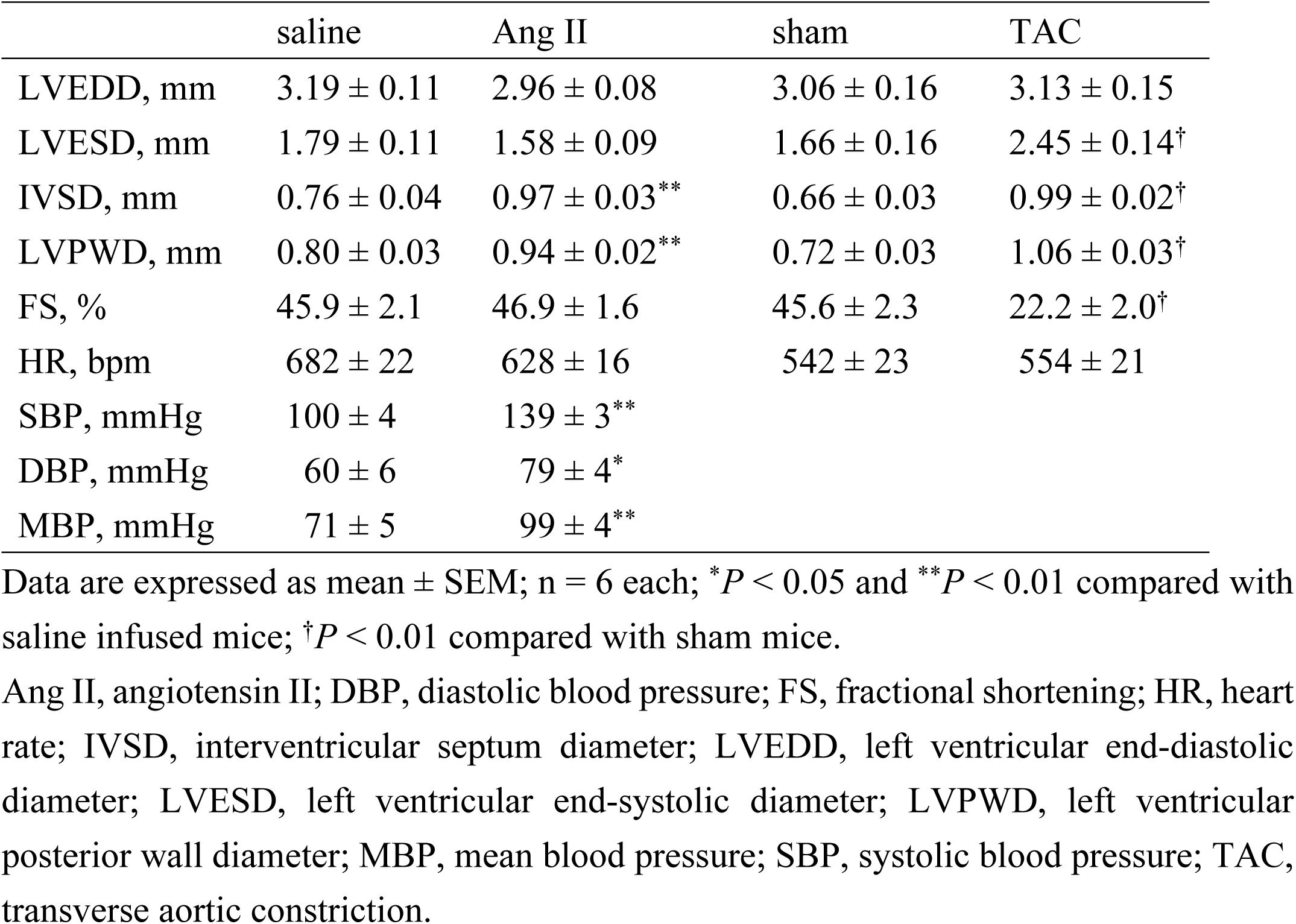
Echocardiographic and hemodynamic data of Ang II infused mice and TAC operated mice.

### Modulation of miR-21 altered PDCD4 expression *in vivo*

MiR-21 has been implicated in fibrosis by suppressing its downstream genes, such as PDCD4, smad family member 7 (Smad7), phosphatase and tensin homolog (PTEN), and sprouty 1 (Spry1) [12, 15, 26, 27]. We examined the mRNA expression levels of these targets using qRT-PCR in Ang II infused mice and TAC mice hearts. PDCD4 mRNA levels were significantly downregulated in Ang II infused mice hearts compared with sham mice. PDCD4 mRNA levels were significantly lower in TAC mice hearts than in sham mice (Fig. 3A). Smad7 mRNA levels were significantly decreased in Ang II infused mice compared with sham mice, although there were no significant differences in PTEN and Spry1 mRNA levels between Ang II infused mice and sham mice (S1A Fig). There were no significant differences in Smad7, PTEN, and Spry1 mRNA levels between TAC mice and sham mice (S1B Fig).

**Fig 3.**
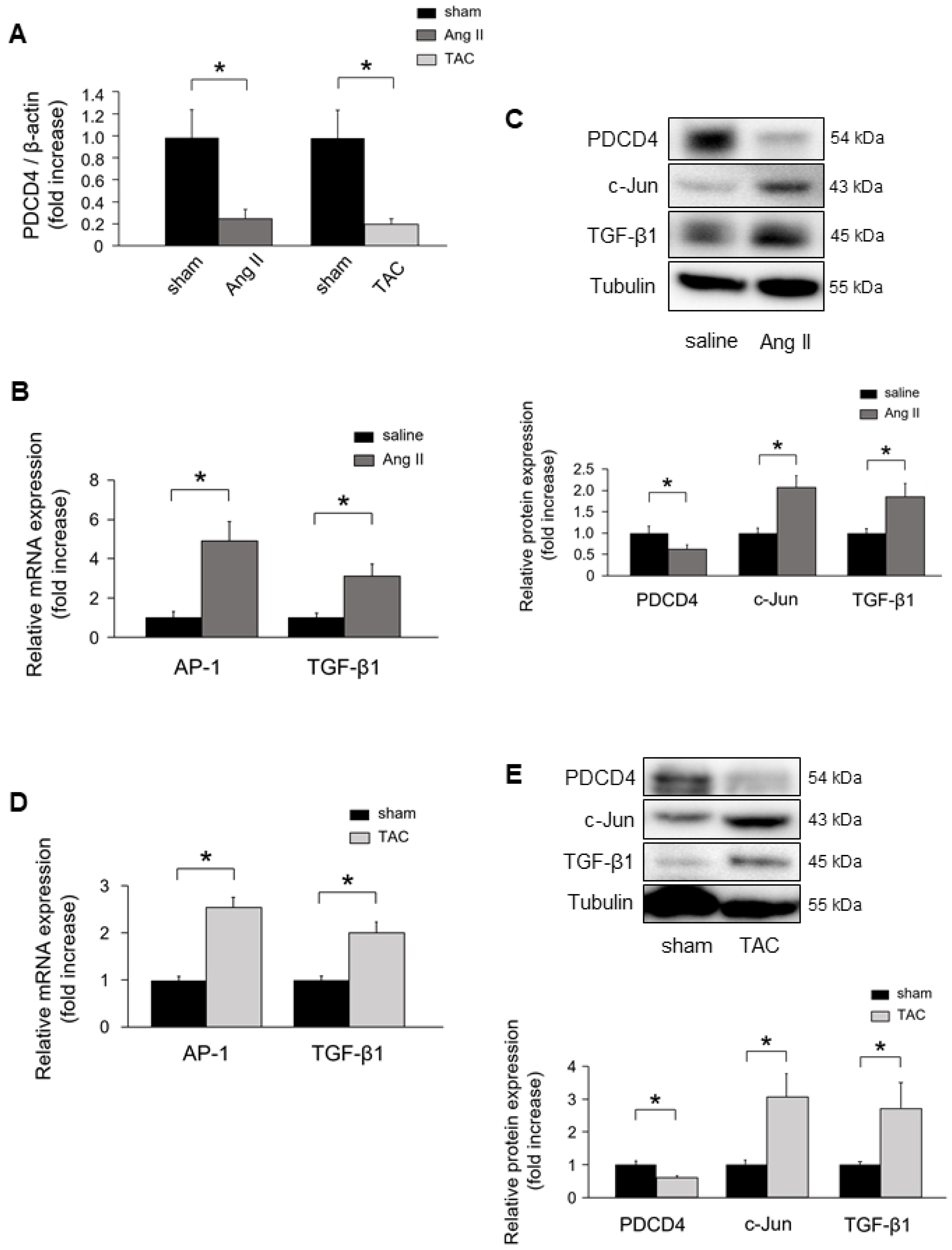
PDCD4 expression and its downstream signaling in Ang II- and TAC-induced cardiac remodeling. **(A)** PDCD4 mRNA expression in Ang II infused and TAC-operated mice (n = 6 per group). **(B)** AP-1 and TGF-β1 mRNA expressions in Ang II infused mice (n = 6 per group). **(C)** Protein expressions of PDCD4, c-Jun, and TGF-β1 in Ang II infused mice (n = 6 per group). **(D)** AP-1 and TGF-β1 mRNA expressions in TAC-operated mice (n = 6 per group). **(E)** Protein expressions of PDCD4, c-Jun, and TGF-β1 in TAC-operated mice (n = 6 per group). Representative images from at least six independent results are shown. Data are expressed as mean ± SEM. \**P* < 0.05.

Since PDCD4 mRNA levels were consistently decreased in Ang II infused mice and TAC mice hearts, we focused on PDCD4. We next investigated PDCD4 downstream target of transcription activator protein 1 (AP-1), a dimeric complex composed of c-Jun and c-Fos family, and TGF-β1 signaling pathway. AP-1 and TGF-β1 mRNA levels were significantly upregulated in Ang II infused mice hearts compared with those of saline infused mice (Fig. 3B). PDCD4 protein expression was significantly decreased in Ang II infused mice hearts, whereas c-Jun and TGF-β1 protein levels were significantly increased compared with those of saline infused mice (Fig. 3C). AP-1 and TGF-β1 mRNA levels were significantly increased in the hearts of TAC mice compared with those of sham mice (Fig. 3D). Moreover, PDCD4 protein expression was significantly decreased in TAC mice hearts, whereas c-Jun and TGF-β1 protein levels were significantly increased compared with those of sham mice (Fig. 3E).

### The impact of miR-21 in cardiomyocytes under hypertrophic stimulation *in vitro*

MiR-21 expression levels were significantly increased in NRCMs under Ang II stimulation (Fig. 4A). PDCD4 mRNA expression was significantly downregulated in NRCMs under Ang II stimulation (Fig. 4B). However, there were no significant differences in mRNA expression levels of Smad7, PTEN, and Spry1 in NRCMs (S2A Fig). The mRNA expression levels of PDCD4, Smad7, and Spry1 were significantly decreased in neonatal rat cardiofibroblasts under Ang II stimulation (S2B Fig). AP-1 and TGF-β1 mRNA levels were significantly upregulated in NRCMs under Ang II stimulation (Fig. 4B). PDCD4 protein expressions were significantly decreased in NRCMs under Ang II stimulation, whereas its targets, c-Jun and TGF-β1 protein expression levels were significantly increased (Fig. 4C).

**Fig 4.**
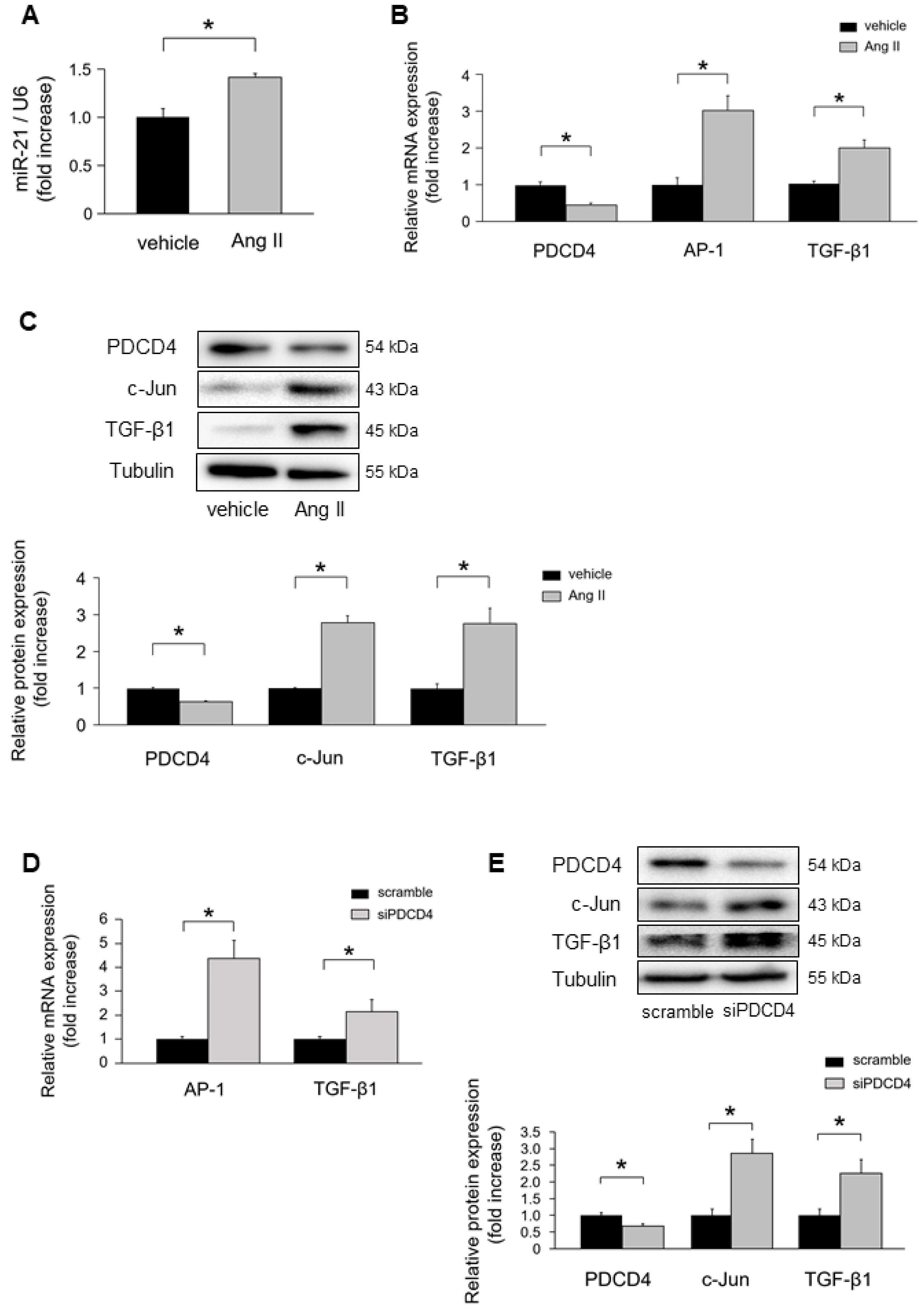
MiR-21 and PDCD4 expressions in cardiomyocytes under Ang II stimulation. **(A)** MiR-21 expressions in cardiomyocytes under Ang II stimulation for 24 h (n = 4–6 per group). **(B)** PDCD4, AP-1, and TGF-β1 mRNA expressions in cardiomyocytes under Ang II stimulation for 24 h (n = 4–6 per group). **(C)** Protein expression levels of PDCD4, c-Jun, and TGF-β1 in cardiomyocytes under Ang II stimulation for 24 h (n = 4–6 per group). **(D)** AP-1 and TGF-β1 mRNA expressions in cardiomyocytes transfected with siPDCD4 (n = 4–6 per group). **(E)** Protein expressions of PDCD4, c-Jun, and TGF-β1 in cardiomyocytes transfected with siPDCD4 (n = 4–6 per group). Representative images from at least four independent results are shown. Data are expressed as mean ± SEM. \**P* < 0.05.

To verify whether PDCD4 directly interacts with AP-1 and subsequent downregulation of TGF-β1, we transfected siPDCD4 into NRCMs. PDCD4 knockdown significantly increased AP-1 and TGF-β1 mRNA levels (Fig. 4D). Western blot analysis also revealed that c-Jun and TGF-β1 protein expression levels were significantly increased by knockdown of PDCD4 (Fig. 4E).

To clarify the direct role of miR-21 in regulating PDCD4 expressions in NRCMs, we transfected mirVana-miR-21-specific inhibitor into NRCMs. MiR-21 inhibitor significantly upregulated PDCD4 mRNA expressions compared with negative control (Fig. 5A). AP-1 and TGF-β1 mRNA expressions were significantly downregulated in NRCMs with miR-21 inhibitor compared with those in the negative control. PDCD4 protein expression levels were significantly increased in NRCMs with miR-21 inhibitor compared with those in the negative control, whereas its targets, c-Jun and TGF-β1 protein expression levels were significantly decreased under miR-21 inhibitor transfection (Fig. 5B).

**Fig 5.**
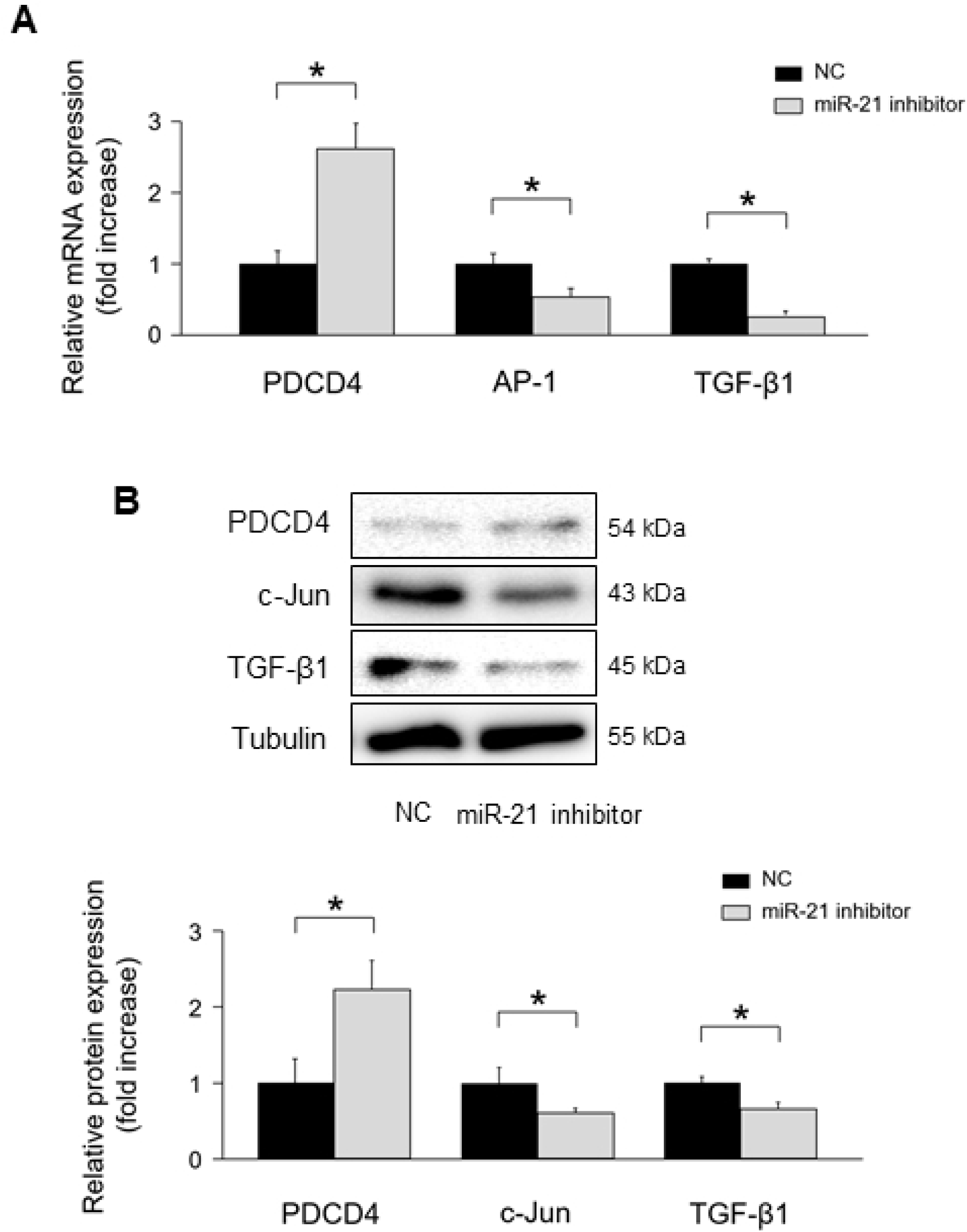
The impact of miR-21 suppression on PDCD4, AP-1, and TGF-β1 signaling in cardiomyocytes. **(A)** Effect of miR-21 suppression on PDCD4, AP-1, and TGF-β1 mRNA expressions in cardiomyocytes (n = 4–6 per group). **(B)** Effect of miR-21 suppression on PDCD4, c-Jun, and TGF-β1 protein expressions in cardiomyocytes (n = 4–6 per group). Representative images from at least four independent results are shown. Data are expressed as mean ± SEM. \**P* < 0.05.

MiR-21 inhibitor attenuated Ang II-induced PDCD4 suppression (Fig. 6A). As a result, subsequent Ang II-induced activation of AP-1 and TGF-β1 mRNA expressions were significantly suppressed by miR-21 inhibitor. PDCD4 protein expression levels were restored by inhibiting miR-21 expressions, whereas c-Jun and TGF-β1 protein expression levels were significantly suppressed (Fig. 6B).

**Fig 6.**
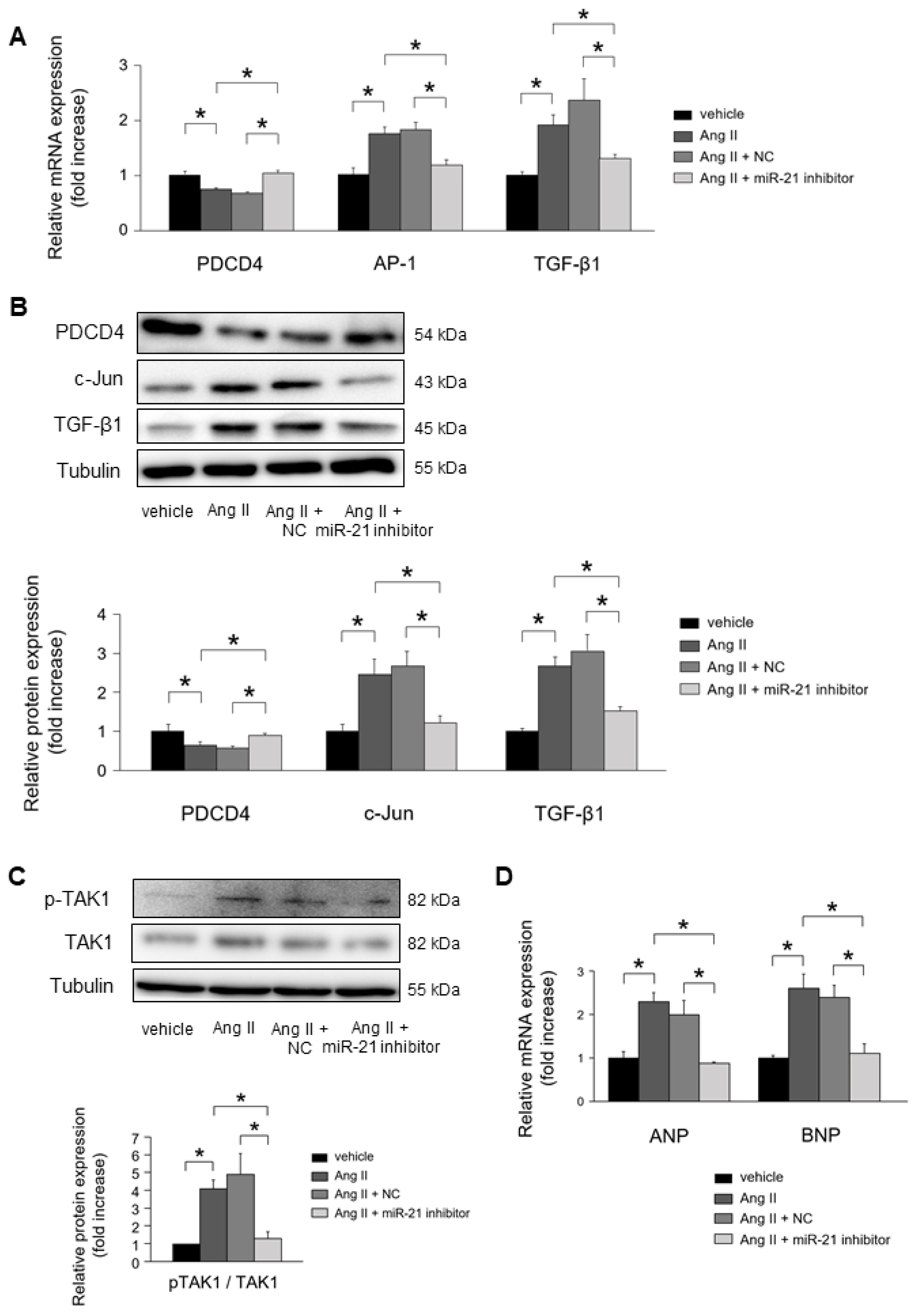
The impact of miR-21 suppression on cardiac remodeling under Ang II stimulation. **(A)** Effect of miR-21 suppression on PDCD4, AP-1, and TGF-β1 mRNA expressions in cardiomyocytes under Ang II stimulation for 24 h (n = 4–6 per group). **(B)** Effect of miR-21 suppression on PDCD4, c-Jun, and TGF-β1 protein expressions in cardiomyocytes under Ang II stimulation for 24 h (n = 4–6 per group). **(C)** Effect of miR-21 suppression on TAK1 and pTAK1 expressions in cardiomyocytes under Ang II stimulation for 24 h (n = 4–6 per group). **(D)** Effect of miR-21 suppression on fetal gene expressions in cardiomyocytes under Ang II stimulation for 24 h (n = 4–6 per group). Representative images from at least four independent results are shown. Data are expressed as mean ± SEM. \**P* < 0.05.

MiR-21 inhibitor significantly suppressed the phosphorylation of TAK1, a key molecular for cardiac hypertrophy, in NRCMs under Ang II stimulation (Fig. 6C). Atrial natriuretic peptide (ANP) and brain natriuretic peptide (BNP) mRNA expressions were significantly upregulated in NRCMs under Ang II stimulation. MiR-21 inhibitor significantly suppressed the mRNA expression of ANP and BNP in NRCMs under Ang II stimulation (Fig. 6D).

## Discussion

In the present study, we revealed the functional role of miR-21 in hypertrophic cardiac remodeling. In patients with HHD, miR-21 expression levels were upregulated in the heart and blood samples. Furthermore, circulating miR-21 levels were positively correlated with serum markers of myocardial fibrosis. In cardiomyocytes, PDCD4 played a pivotal role in regulating cardiac remodeling as a target gene of miR-21 under hypertrophic stimulation. Knockdown of miR-21 ameliorated AP-1 mediated TGF-β1 signaling through regulating PDCD4 in cardiomyocytes. To the best of our knowledge, this study is the first report to evaluate the impact of miR-21 in patients with HHD, and that in cardiomyocytes in hypertrophic cardiac remodeling using *in vivo* and *in vitro* experiments. A schema that includes the suggested pathway from the present study is shown in Fig. 7.

**Fig 7.**
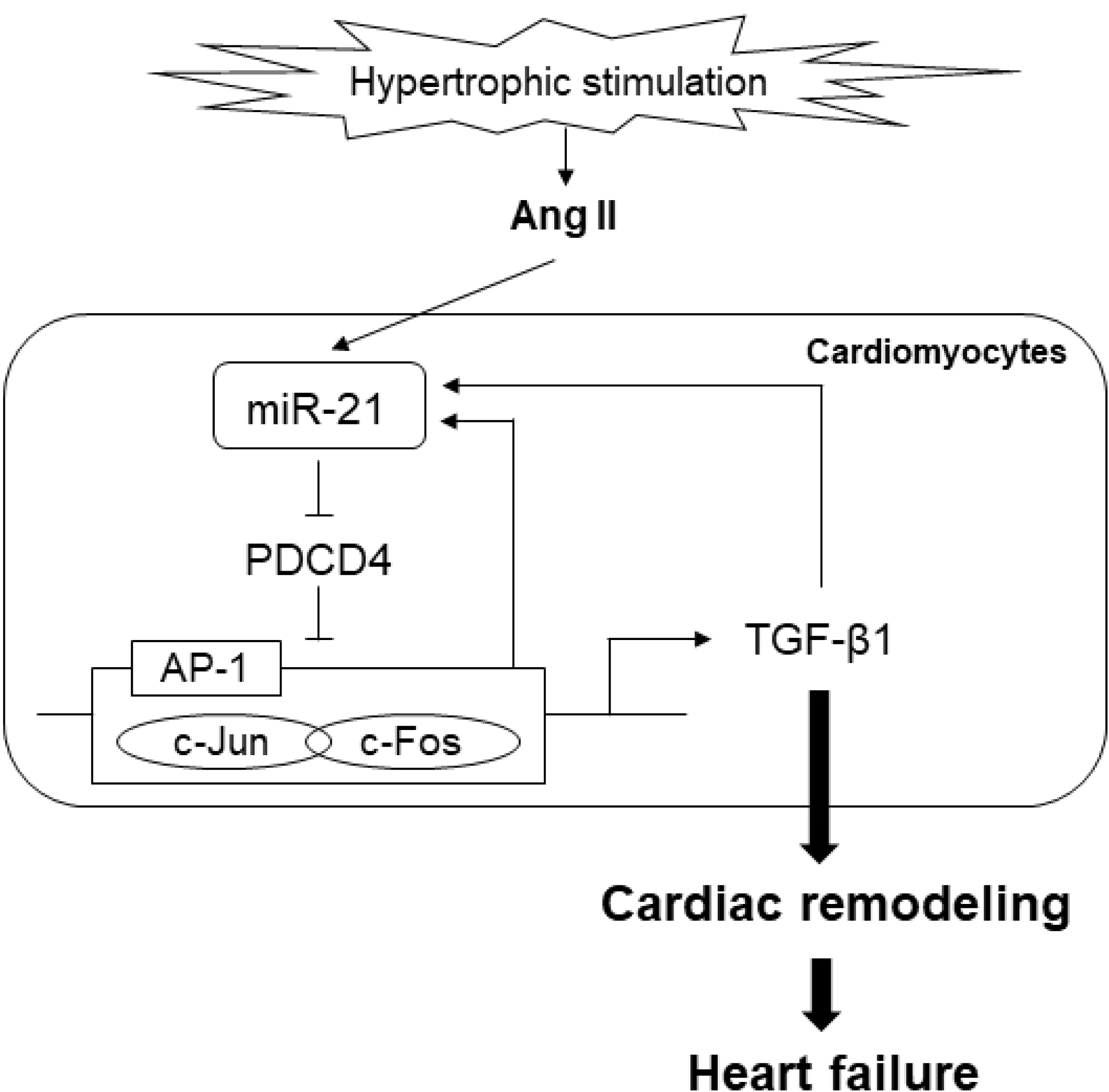
A schema that includes the proposed pathway of miR-21 in cardiac remodeling.

The pathogenetic mechanism of HHD is thought to be related to cardiac remodeling, including cardiac fibrosis and hypertrophy [28]. Hence, regression of hypertrophic cardiac remodeling can improve the prognosis of patients with hypertension. The contribution of miRs in the cardiomyopathies such as ischemic heart disease, hypertrophic cardiomyopathy, and dilated cardiomyopathy has been shown [29–31]. However, the impact of miR-21 as well as other miRs on the pathogenesis of HHD, which is one of the most important hypertension-induced organ damages is still unclear. In the present study, we showed that miR-21 expression levels were significantly increased in both serum and heart samples of patients with HHD compared with normal subjects. Moreover, there were significant positive correlations between circulating miR-21 levels and serum markers of myocardial fibrosis. These findings support the association between miR-21 and cardiac remodeling in patients with HHD.

It is well known that hypertension-derived mechanical stress induces Ang II synthesis, and subsequent activation of nuclear AP-1 leads to the upregulation of TGF-β1 [32]. TGF-β1 induces fibroblast-to-myofibroblast differentiation and extracellular matrix (ECM) production, which leads to cardiac fibrosis [33]. It was reported that TGF-β1 was increased with advancing of fibrosis in the hearts of patients who underwent cardiac surgery [34]. While mice with systemic overexpression of TGF-β1 showed cardiac fibrosis and hypertrophy, mice with systemic knock-out of TGF-β1 ameliorated Ang II-induced cardiac hypertrophy [35, 36]. TGF-β1 is a key mediator of the pathogenesis of cardiac remodeling under hypertrophic stimulation. Interestingly, activation of TGF-β1 signaling increases miR-21 expressions [13]. In contrast, several reports revealed that miR-21 can activate TGF-β1 signaling [11, 12]. This interrelationship forms interesting positive feedback loop. Our results *in vivo* study showed that miR-21 and TGF-β1 expression levels were significantly increased in Ang II infused mice and TAC mice, suggesting that an interrelationship between miR-21 and TGF-β1 may play an important role in hypertrophic cardiac remodeling.

Elevated miR-21 expression levels were reported to be associated with organ fibrosis, such as lung, kidney, liver, and heart via promoting fibroblast activation [12, 16, 37, 38]. Several reports have shown the fibrogenic function of miR-21 in fibroblasts through modulation of its target genes, such as PDCD4, Smad7, PTEN, and Spry1 [12, 15, 26, 27]. Thus, although the functional role of miR-21 in fibroblast is well known, there were few studies assessing the functional role of miR-21 in cardiomyocyte. In the present study, we found that Ang II significantly upregulated PDCD4 mRNA expression in cardiomyocytes, although there were no significant differences in the mRNA expression levels of Smad7, PTEN, and Spry1. These results suggest that PDCD4 plays an important role in regulating cardiac remodeling as a target gene of miR-21 under hypertrophic stimulation in cardiomyocyte.

PDCD4 is a well-known tumor suppressor and is involved in apoptosis. It was reported to be a powerful inhibitor of AP-1 [39]. On the other hand, activation of AP-1 upregulates miR-21 expressions [40]. In the present study, we showed that PDCD4 was significantly decreased and AP-1 was increased in Ang II infused mice and TAC mice. In addition, AP-1 mediated TGF-β1 expression was significantly upregulated under Ang II stimulation *in vitro*. Thus, there arises a possibility that miR-21 might enhance its own transcription through miR-21/PDCD4/AP-1 pathway and exacerbate the fibrogenic process in hypertrophic cardiac remodeling.

Cardiomyocytes and fibroblasts cooperatively regulate cardiac cell signaling via paracrine mediators, which is involved in cardiac remodeling [41]. It has been reported that TGF-β1 was induced in response to hypertrophic stimuli not only in fibroblasts but also in cardiomyocytes, and acting in a paracrine and/or autocrine manner [42, 43]. However, although the effects of miR-21 inhibition on cardiac remodeling were demonstrated in fibroblast, its effects in cardiomyocyte were poorly understood. In the present study, knockdown of miR-21 expression rescued Ang II-induced PDCD4 suppression. Furthermore, knockdown of miR-21 significantly suppressed Ang II-induced AP-1 and TGF-β1 signaling in cardiomyocytes. These results suggest that inhibition of miR-21 prevents hypertrophic stimulation-induced cardiac fibrosis by suppressing miR-21/PDCD4/AP-1 feedback loop.

In addition to fibrogenic function of TGF-β1, Koitabashi et al. showed that suppression of myocyte-derived TGF-β1 ameliorated cardiac hypertrophy by inhibiting non-canonical pathways, in particular TAK1 [44]. Consistently, we showed that Ang II stimulation induced TAK1 activation in cardiomyocytes. Furthermore, we showed that inhibition of miR-21 expression suppressed TAK1 activity and subsequent fetal gene expressions in cardiomyocytes. Remarkably, Thum et al. demonstrated that silencing of miR-21 *in vivo* attenuated cardiac fibrosis and hypertrophy under pressure overload stimulation through deactivation of cardiac fibroblast [15]. Taking our results into consideration, this beneficial effect of miR-21 inhibitor in suppressing hypertrophic cardiac remodeling might be attributed to not only cardiac fibroblast but also cardiomyocyte.

We need to point out several limitations of our study. First, 3 patients with hypertension were included in control group, although they had normal cardiac function and had no left ventricular hypertrophy. Second, because there were 6 patients with diabetes mellitus in HHD group, we could not completely rule out the possibility of the influence of diabetes mellitus on cardiac remodeling. Third, EMB study size was relatively small for investigating the impact of miR-21 on cardiac remodeling in patients with HHD. Finally, we have not evaluated the effect of miR-21 inhibitor *in vivo*. Several studies demonstrated that inhibition of miR-21 suppressed cardiac remodeling by regulating cardiac fibroblast [15, 16, 26], although we confirmed the protective effect of miR-21 inhibitor in cardiomyocyte under hypertrophic stimulation *in vitro*.

## Conclusions

MiR-21 was associated with fibrogenesis in heart under hypertrophic stimulation. Inhibition of miR-21 expressions prevent hypertrophic cardiac remodeling by regulating PDCD4 and AP-1, TGF-β1 signaling pathway.

## Acknowledgement

We thank Ms. Emiko Nishidate and Ms. Hiroko Sasaki for their excellent technical assistance and comments.

## Supporting information

**S1 Fig. Smad7, PTEN, and Spry1 mRNA levels in Ang II infused and TAC mice.**

**(A)** Smad7, PTEN, and Spry1 mRNA levels in Ang II infused mice compared with those of saline infused mice (n = 6 per group). **(B)** Smad7, PTEN, and Spry1 mRNA levels in TAC mice compared with those of sham mice (n = 6 per group). Data are expressed as mean ± SEM. \**P* < 0.05.

**S2 Fig. The differences in the expression levels of miR-21 targets between cardiomyocytes and cardiofibroblasts.**

**(A)** The mRNA expressions of Smad7, PTEN, and Spry1 after treatment with vehicle or Ang II for 24 h in NRCMs (n = 4–6 per group). **(B)** The mRNA expressions of PDCD4, Smad7, PTEN, and Spry1 after treatment with vehicle or Ang II for 24 h in cardiofibroblasts (n = 4–6 per group). Data are expressed as mean ± SEM. \**P* < 0.05.

